# Adolescent Stress Confers Resilience to Traumatic Stress Later in Life: Role of the Prefrontal Cortex

**DOI:** 10.1101/2021.03.16.435691

**Authors:** E.M. Cotella, N Nawreen, R.D. Moloney, S.E. Martelle, K.M. Oshima, P. Lemen, J. N. NiBlack, R. R. Julakanti, M. Fitzgerald, M.L. Baccei, J.P. Herman

## Abstract

Stress during adolescence is usually associated with psychopathology later in life. However, under certain circumstances, developmental stress can promote an adaptive phenotype, allowing individuals to cope better with adverse situations in adulthood, thereby contributing to resilience. The aim of the study was to understand how adolescent stress alters behavioral and physiological responses to traumatic stress in adulthood. Sprague Dawley rats were subjected to adolescent chronic variable stress (adol CVS) followed by single prolonged stress (SPS) in adulthood. One week after SPS, animals were tested for acquisition, extinction, extinction recall and reinstatement of auditory-cued fear conditioning, with neuronal recruitment during reinstatement assessed by Fos expression. Patch clamp electrophysiology was performed to examine physiological changes associated with resilience. We observed that adol CVS blocked SPS-induced impairment of extinction learning (males) and enhancement fear reinstatement (both sexes). SPS effects were associated with a reduction of infralimbic (IL) cortex neuronal recruitment after reinstatement in males and increased engagement of the central amygdala in females, both of which were also prevented by adol CVS. We explored the mechanism behind reduced IL recruitment in male rats by studying the intrinsic excitability of IL pyramidal neurons. SPS reduced excitability of IL neurons and prior adol CVS prevented this effect, indicating that adolescent stress can impart resilience to the effects of traumatic stress by modification of IL output in males. Overall, our study suggests that prior stress exposure can limit the impact of a subsequent severe stress exposure in adulthood, effects that are mediated by sex-specific modification of infralimbic and amygdala signaling.

## Introduction

Understanding factors that affect the brain during adolescence has substantial health relevance, given the onset of numerous affective conditions during this developmental period (e.g., depression, anxiety disorders) (Andersen and Teicher, 2008; Kessler et al., 2005; Paus et al., 2008). In general, chronic stress during development is associated with the emergence of pathology, particularly when occurring during early life (de Kloet et al., 2005; Heim et al., 2008; Oitzl et al., 2010; Riboni and Belzung, 2017; Tost et al., 2015). However, mild to moderate stress during some development periods may also promote an adaptive response to adverse situations later in life, contributing to stress resilience (Ordoñes Sanchez et al., 2021; Ricon et al., 2012; Romeo, 2015; Schmidt, 2011; Southwick and Charney, 2012). Previous work from our lab indictates that chronic variable stress (CVS) during adolescence can evoke specific effects later in life that may determine either risk or resilience (Cotella et al., 2020, 2019). While the mechanisms implicated in developmental vulnerability to stress dysregulation are widely studied, resiliency after stress is poorly understood.

Memories acquired under stressful situations are usually strongly consolidated and can be retrieved more easily than those acquired in neutral situations (Meir Drexler and Wolf, 2017). Prior exposure to stress can further enhance the acquisition and/or expression of the fear related behaviors (Blouin et al., 2016), processes linked to the prelimbic (PL) and infralimbic (IL) divisions of the rodent medial prefrontal cortex (Giustino and Maren, 2015). Learned fear has an obvious adaptive value, increasing the chance of survival in life threatening situations (Giustino and Maren, 2015). However, traumatic experiences can lead to exaggerated and prolonged fear responses that can have pathological consequences, as seen in post-traumatic stress disorder (PTSD). Here, individuals experience recurring episodes of involuntary memories associated with an intense stress response, resulting in hyperalertness and avoidance of situations that remind them of the traumatic event (Blouin et al., 2016; Sareen, 2014). Interestingly, although there is a high chance of experiencing trauma in the population, only about 7% of people develop PTSD (Benjet et al., 2016; Kessler et al., 2005), suggesting that resilience or vulnerability to development of PTSD may be determined by experiential and/or genetic factors.

Rodents are widely used to study how stress affects learned fear memories. Stress-enhanced fear models usually combine exposure to one or more stressors, with fear responses tested in a conditioning paradigm (Blouin et al., 2016). One of the most widely-used and reproduced models is the single-prolonged stress protocol (SPS) developed by Liberzon (Liberzon et al., 1999, 1997). Exposure to SPS impairs extinction and extinction recall of a fear conditioned response one week later (Knox et al., 2012b, 2012a; Kohda et al., 2007), comprising a late-emerging enhancement of fear, as is characteristic of PTSD.

Prefrontal activity and neuronal intrinsic excitability is associated with stress resilience and vulnerability (Kumar et al., 2014). For example, in humans the aberrent fear response in PTSD is associated with ventromedial PFC (homolog to the rodent IL)(Öngür and Price, 2000) hypoactivity and loss of top-down control over the amygdala (Milad et al., 2009). In rodents, SPS also reduces neuronal activation in the IL (Piggott et al., 2019), which may play a role in the abnormal fear extinction deficits associated with SPS. Conversely, optogenetic drive of the mPFC can promote stress resilience, and successful stress coping is linked to elevated mPFC activation after social defeat stress (Covington et al., 2010) . However, the circuitry underlying vulnerability and resilience are largely unknown (Russo et al., 2012).

In the present study we assess the impact of adolescent CVS on stress vulnerability or resilience to subsequent SPS in adulthood. Our data indicate that the experience of stress during adolescence blocks fear potentiation following SPS, leading to resilience, a phenomenon that can be linked to descreases in intrinsic excitability of IL mPFC glutamatergic pyramidal neurons.

## Results

### Experiment 1: Cued conditioned response

Fig. 1.C-F illustrates the conditioned freezing response throughout the different sessions of the fear conditioning paradigm in animals that were submitted to chronic variable stress during adolescence (adol CVS) and later subjected to single prolonged stress (SPS) in adulthood. Animals were submitted to a tone-conditioned paradigm as shown in Fig 1.B. Fig 1.C-D show the effects for each phase of paradigm evaluated.

**Figure 1:**
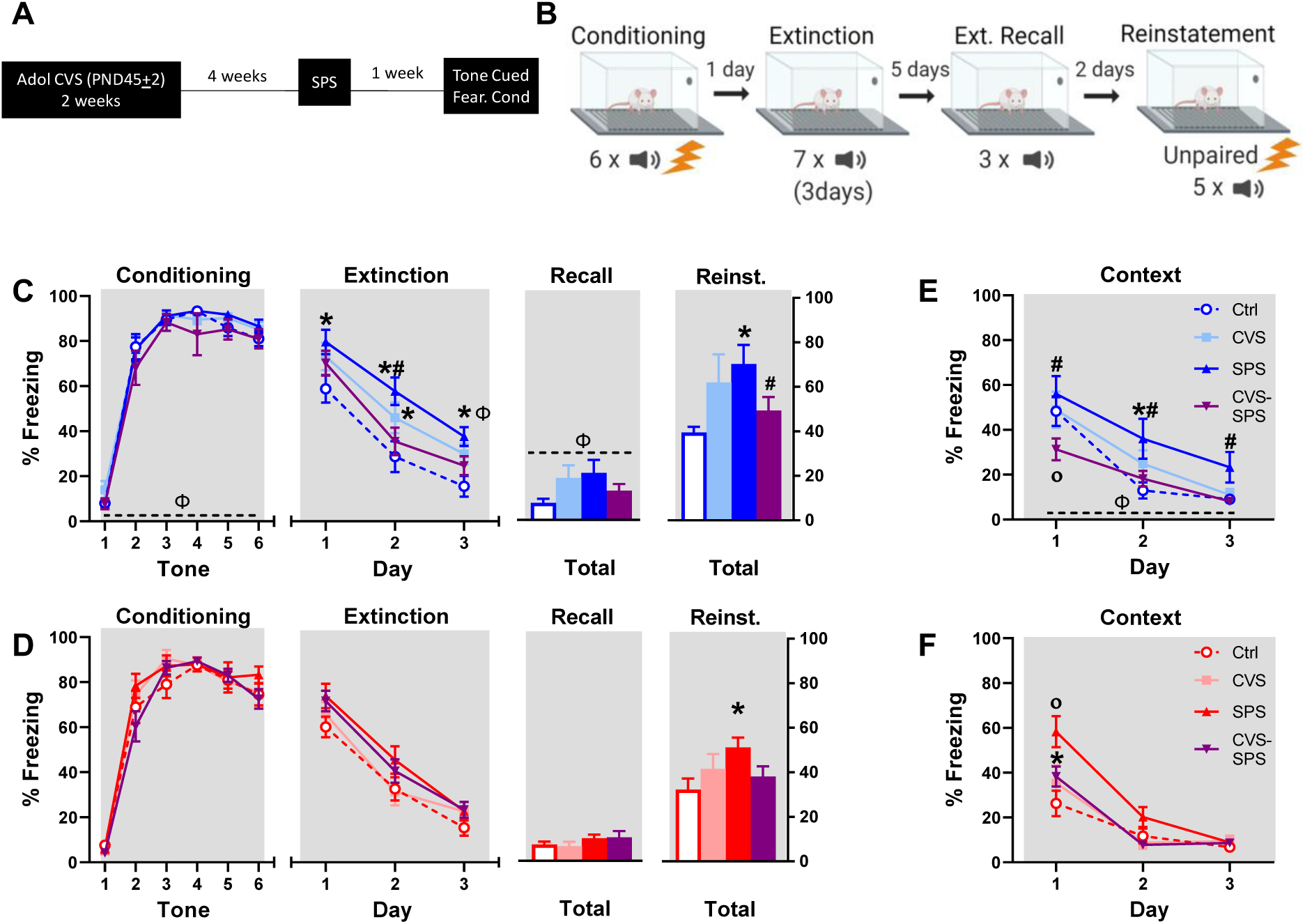
**A)** Experimental timeline. **B)** Fear conditioning paradigm. **C, D)** Previous adolescent stress prevented SPS extinction deficit in male rats and prevented reinstatement of tone-conditioned freezing after SPS in both male and female rats. Multiple planned comparisons: *p<0.05 compared to same sex control group, # p<0.05 compared to same sex adol CVS-SPS (extinction) or SPS (reinstatement) group. □ = sex effect. **E-F)** Context conditioning was tested as freezing during the 2 minutes prior to the first tone each day of the extinction procedure. Previous adolescent stress reduced conditioned response to context in SPS animals re exposure (day 1) in both sexes and enhanced extinction of this response in male rats. Male rats had higher freezing than females (p<0.05 □). Individual planned comparisons: *p<0.05 compared to same sex control group, # p<0.05 compared to same sex adol CVS-SPS, ° p<0.05 compared to all other same sex groups. Data represented as Mean±SEM. Image created with BioRender.com

### Conditioning

None of the treatments had effects on the conditioning phase. There was an interaction between adol CVS x SPS (F_(1,79)_= 5.075, p=0.027) but no individual differences in the Bonferroni test. The significant effect of time (F_(5,395)_= 469.308, p<0.0001) confirmed conditioning of the response. There was main effect of sex (F_(1,79)_= 7.724, p=0.007), with Bonferroni comparisons indicating a general higher expression of freezing in male rats.

### 3-day extinction

There were significant effects of SPS (F_(1,76)_= 6.698, p=0.012) and time (F_(2,152)_= 475,661, p<0.0001) and an adol CVS x SPS interaction (F_(1,76)_= 7.414, p= 0.008), with no effect of sex. The CVS x SPS post hoc analysis confirmed that, in general, SPS groups had higher freezing levels compared to controls over the whole extinction procedure regardless of sex. When performing planned comparisons by sex, only male rats showed statistical significant effects, with the SPS group having higher freezing than the control group on all three days (p<0.05) (the CVS only group had enhanced freezing only on day 2 (p<0.05)). Prior adol CVS prevented the SPS effects, as the double-hit group remained at control levels on all testing days and had significantly less freezing than the SPS group on day 2 (p<0.05). Sex differences were observed only on day 3, with SPS evoking higher freezing in male rats (p<0.05).

### Recall

The levels of extinction attained were stable for both sexes as tested in the recall phase. In this case, five days after extinction, animals received a brief extra extinction session (3 tones) to test for possible spontaneous recovery of the conditioned response and to corroborate that the levels of freezing in all the groups were equal before reinstatement. We observed a main effect of sex (F_(1,76)_=7.958, p=0.006) with males expressing more freezing in general, and a triple interaction sex x adol CVS x SPS (F_(1,76)_=4.792, p=0.032) with no group differences emerging for any individual Bonferroni comparison.

### Reinstatement

We observed a significant adol CVS x SPS interaction F_(1,76)_= 11.8095, p=0.001. Posthoc comparison indicated that regardless of sex, the SPS group expressed higher freezing than the control group (p<0.05 respectively), while the double-hit group prevented the effect of SPS, remaining at control freezing levels and expressing significantly less freezing time than the SPS group (p<0.05). When analyzing the individual responses by sex (planned comparisons), we observed that in female rats, only the SPS group differed from control (p<0.05). In the case of male rats, the SPS group had higher freezing than control (p<0.05) and the adol CVS-SPS group (p<0.05).

### Context conditioned response

To quantify the conditioned response to the conditioning context, we evaluated the freezing response evoked every day of the extinction procedure before the first tone was presented and analyzed the progression over 3 days (Fig 1.E-F).  We observed a main effect of adol CVS F_(1,76)_=4.097, p=0.046 and significant adol CVS x SPS interaction F_(1,76)_=11.729, p=0.001. The posthoc analysis indicated that animals subjected to SPS expressed higher freezing when re-exposed to the conditioning context compared to all the other groups (p<0.05 respectively). There was a sex effect F_(1,76)_=6.971, p=0.01, with male rats having more freezing time. There was also an effect of time F_(2,152)_=172.946, p<0.00001, indicating reduction of freezing to subsequent exposure. Finally, there was a time x sex x SPS (F_(2,152)_=9.253, p=0.0002) interaction. Planned comparisons showed that male rats subjected to SPS alone expressed higher freezing than the control group on day 2 while the group subjected to the double-hit model of stress had less context freezing compared to all the other groups on day 1 (p<0.05 respectively). This difference was maintained against the SPS group on the rest of the days tested (P<0.05 respectively). In the case of females, SPS group had higher freezing to the context than all the other groups on day 1 (p<0.05 respectively) and the adol CVS-SPS group also had more freezing compared to controls on that day (p<0.05).

### Fos expression after reinstatement

Figure 2 summarizes Fos activation (Fos immunoreactivity, Fos-ir) in the mPFC and amygdala assessed following the reinstatement trial (90 min after the onset of the session). In the case of males, we observed a significant effect of SPS in the infralimbic (IL) cortex of male rats, F_(1, 16)_=7.706, p=0.0135. Planned comparisons confirmed that the SPS groups had significantly less Fos-ir than the control and CVS animals (p<0.05), while the other groups did not differ from controls (Fig 2.A). In females, effects of adol CVS and SPS were only observed in the central nucleus of the amygdala, particularly, in lateral (CeL) but not medial subdivision of the central amygdala (CeM) (SPS F_(1, 23)_= 5.945, p=0.0229, adol CVS x SPS F _(1, 23)_ = 7.241, p=0.0130). Bonferroni’s test confirmed that SPS significantly increased neuronal recruitment (p<0.05) and this was prevented in the double-hit group (p<0.05) (Fig 2.B).

**Figure 2:**
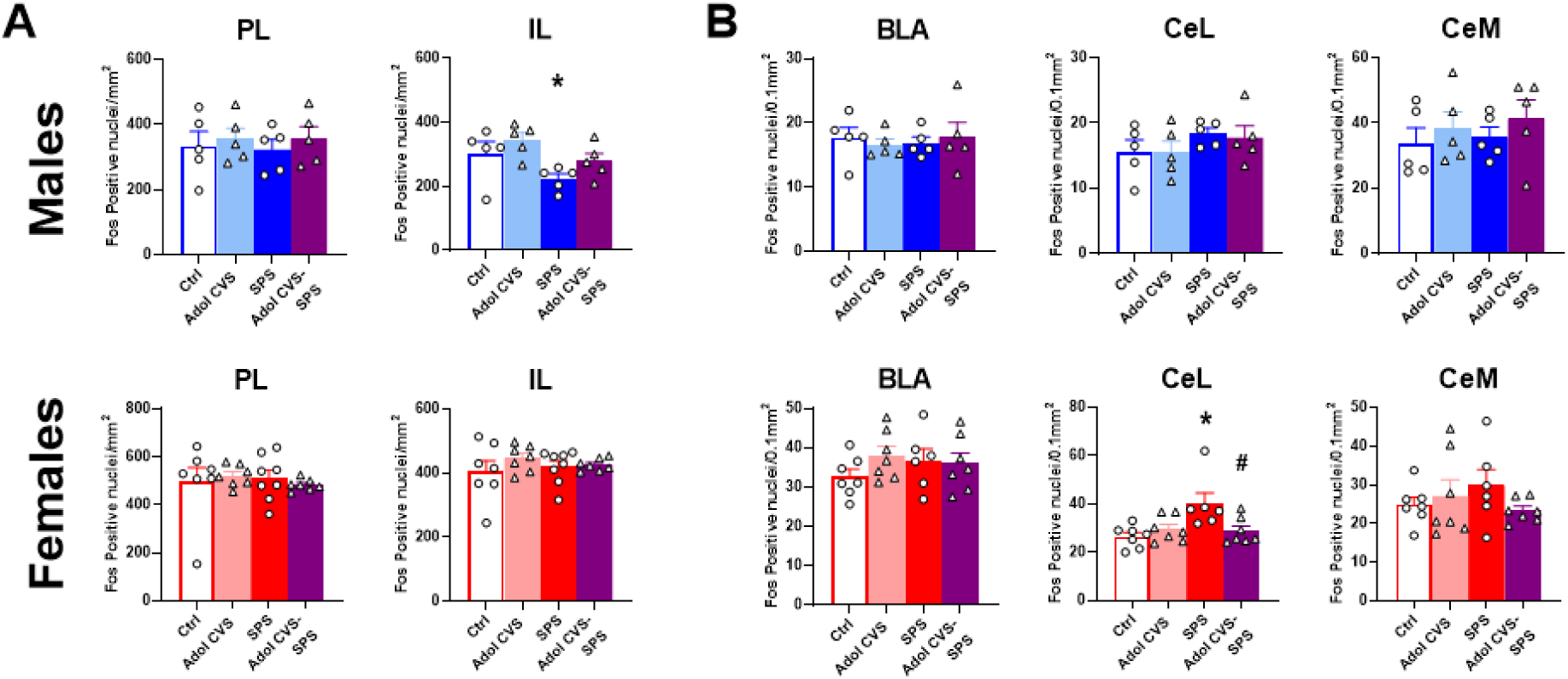
Fos immunoreactivity (Fos-ir) in the medial prefrontal cortex **(A)** and amygdala **(B)** of the animals 90 min after reinstatement (see timeline figure 1.B). There was significant effect of SPS in the infralimbic division of male rats. Planned comparisons confirmed less Fos-ir in the SPS compared to control animals (p<0.05) (*). None of the other groups differed from controls. In females, effects of adol CVS and SPS were only observed in the central nucleus of the amygdala, particularly, in lateral portion (CeL) and not on the central medial division (CeM). Bonferroni’s test confirmed that SPS significantly increased neuronal recruitment (p<0.05 *) and this was prevented in the double-hit group (p<0.05 #). Data as Mean+SEM.

### Experiment 3: Electrophysiology

We next investigated the potential cellular mechanisms underlying how adolescent stress can prevent SPS-induced changes in fear behavior and Fos activation in IL in male rats. Male rats were selected based on the clear effects of both SPS and adolCVS/SPS on extinction learning, a process linked to the IL. We measured the intrinsic membrane properties and firing frequency of IL pyramidal neurons in layer V, the major source of subcortical output from the IL (Baker et al., 2018). We found that prior experience of adol CVS prevented SPS-mediated changes in intrinsic excitability of IL pyramidal neurons. There was a significant main effect of SPS (F_(1,99)_= 32.3, p<0.0001^a (table 1)^) and adol CVS (F_(1,99)_= 8.5, p=0.005^a^) on rheobase (Fig.3D). SPS significantly increased rheobase compared to the control group (p<0.05), which was prevented by prior adol CVS (p<0.05 compared to SPS). There was a significant main effect of SPS (F_(1,98)_= 41.96, p<0.0001^b^), adol CVS (F_(1,98)_= 21.7, p=0.0002^b^) and a significant adol CVS X SPS interaction (F_(1,98)_= 6.7, p=0.01^b^) on membrane resistance (Fig.3E). Bonferroni’s test indicated that SPS significantly decreased membrane resistance (p<0.05), which was prevented by prior adol CVS (p<0.05 compared to SPS). Statistical analysis for membrane resistance was performed on log transformed data. There were significant main effects of SPS (F_(1,92)_=19, p<0.0001^c^), adol CVS (F_(1,92)_=14.3, p=0.0003^c^) and a significant adol CVS x SPS interaction (F_(1,92)_=5.0, p=0.02^c^) on action potential (AP) threshold (Fig 3F) . Bonferroni’s test indicated that SPS significantly increased AP threshold compared to controls (p<0.05), with prior adol CVS preventing the effect (p<0.05 compared to SPS). There were main effects of SPS (F_(1,97)_=20, p<0.001^d^), adol CVS (F_(1,97)_=25.9, p<0.0001^d^) and significant adol CVS x SPS interaction (F_(1,97)_= 6.5, p=0.01^d^) on action potential amplitude (Fig.3G). Bonferroni’s test indicated that SPS significantly lowered AP amplitude compared to controls (p<0.05), and prior experience adol CVS prevented it (p<0.05 compared to SPS). There was a significant effect of SPS (F_(1,97_)=8.3, p=0.005^e^) and adol CVS (F_(1,97)_= 42.2, p<0.0001^e^) on AP50 (Fig. 3H). Planned comparisons indicated a decrease in AP50 following SPS compared to control (p<0.05) and prior adol CVS prevented that effect (p<0.05 compared to SPS). Increase in AP50 was also observed following adol CVS only (p<0.05 compared to control). Resting membrane potential (RMP) was unaltered among the groups (Fig.3I). 2 way ANOVA revealed no significant main effect of SPS (F_(1,95)_=1.3, p=0.3), adol CVS (F_(1,95)_=0.02, p=0.9) or SPSx adol CVS interaction on RMP (F_(1,95)_=0.1, p=0.7).

**Figure 3:**
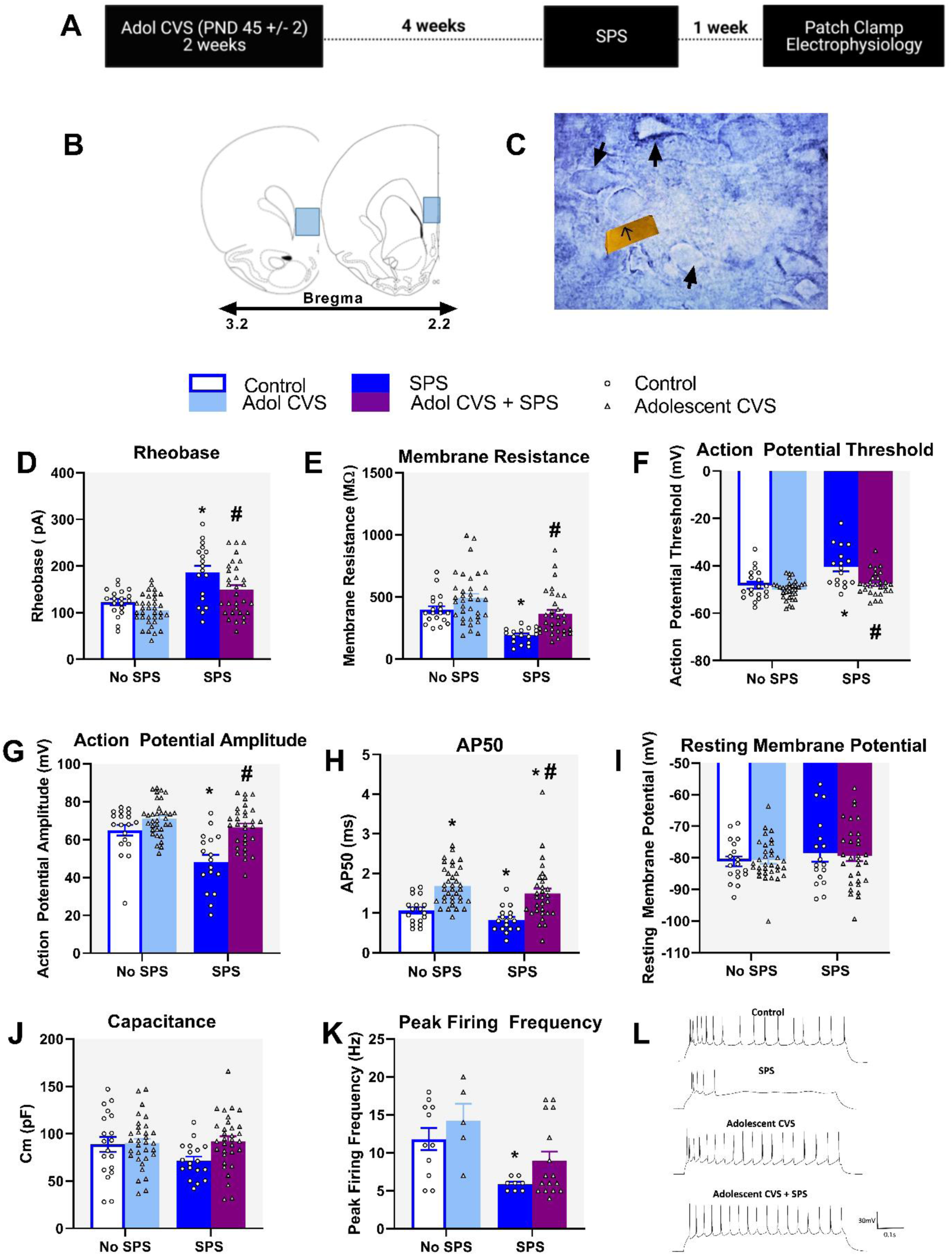
Adolescent CVS prevents SPS effects on membrane properties and intrinsic excitability of IL pyramidal neurons. **(A)** Experimental timeline; **(B)** Schematic of coronal brain sections through PFC where recordings were performed, blue boxes indicate infralimbic region of the PFC; **(C)** Pyramidal neurons were identified based on somal morphology and presence of prominent apical dendrite . Arrows indicate pyramidal neurons; SPS increased rheobase **(D)** and decreased membrane resistance **(E)**, whereas prior experience of adol CVS was able to prevent these effects. SPS increased the threshold for action potential (AP) firing **(F)** and decreased AP amplitude **(G)**, both of which were prevented by prior adol CVS. SPS also reduced the duration of AP (AP 50), which was also blocked by prior adol CVS. It should be noted that adol CVS alone increased AP duration **(H)**. Finally, adol CVS was also able to attenuate the reduction in peak firing frequency observed following SPS **(K)**. No changes in resting membrane potential **(I)** or membrane capacitance **(J)** were observed. **(L)** demonstrates representative traces of action potentials evoked by 20pA current injection for the respective groups; For D,H,K * and # represents planned comparison effects compared with control and SPS respectively. For E, F and G* and # represents post hoc Bonferroni effects compared with control and SPS respectively. Data represented as Mean±SEM.

Membrane capacitance was unaltered among groups (Fig.3J) indicating the treatments did not likely affect cell size. 2 way ANOVA of membrane capacitance revealed no significant main effect of SPS (F_(1,97)=_1.8, p=0.2), Adol CVS (F_(1,97)=_3.6, p=0.06) or SPSx adol CVS interaction (F_(1,97)=_2.9, p=0.09). Analysis of peak firing frequency revealed a significant main effect of SPS (F_(1,36)_=13.4, p=0.0008^h^). Planned comparisons indicated that SPS significantly reduced peak firing frequency compared to controls (p<0.05), whereas the prior adol CVS+SPS group did not differ from the control group (Fig.3K). Figure 3L shows representative traces of action potentials evoked by 20pA current injection for the respective groups. Together these data indicate that prior experience of adolescent stress is able to prevent the reduction in intrinsic excitability and firing rate of IL layer V pyramidal neurons following SPS.

## Discussion

Our results strongly suggest that prior experience with stress during adolescence can evoke a resilient phenotype in the adult, characterized by the prevention of the effects of SPS in a fear conditioning paradigm and on IL pyramidal cell excitability. Our data indicate that the adaptations resulting from exposure to chronic stress during adolescence buffer the behavioral impact of a model of traumatic stress in adulthood, blocking known effects of SPS on subsequent fear potentiation.

While prior CVS is able to block enhancement of reinstatement in both sexes, it appears to do so by distinct neuronal mechanisms. Reversal of SPS-induced resinstatement was accompanied by IL hypoactivity in males and CeL recruitment in females, suggestion differential engagement of cortical regions regulating extinction (males) vs. fear expression (females) across the sexes. Notably extinction deficits were only observed in males, consistent with the known role of the IL in extinction of conditioned fear. A role for the IL in CVS-induced resilience in males is further supported by hypoactivity of layer V pyramidal cells following SPS, which is blocked by prior adolescent CVS (Figure 4).

**Fig 4:**
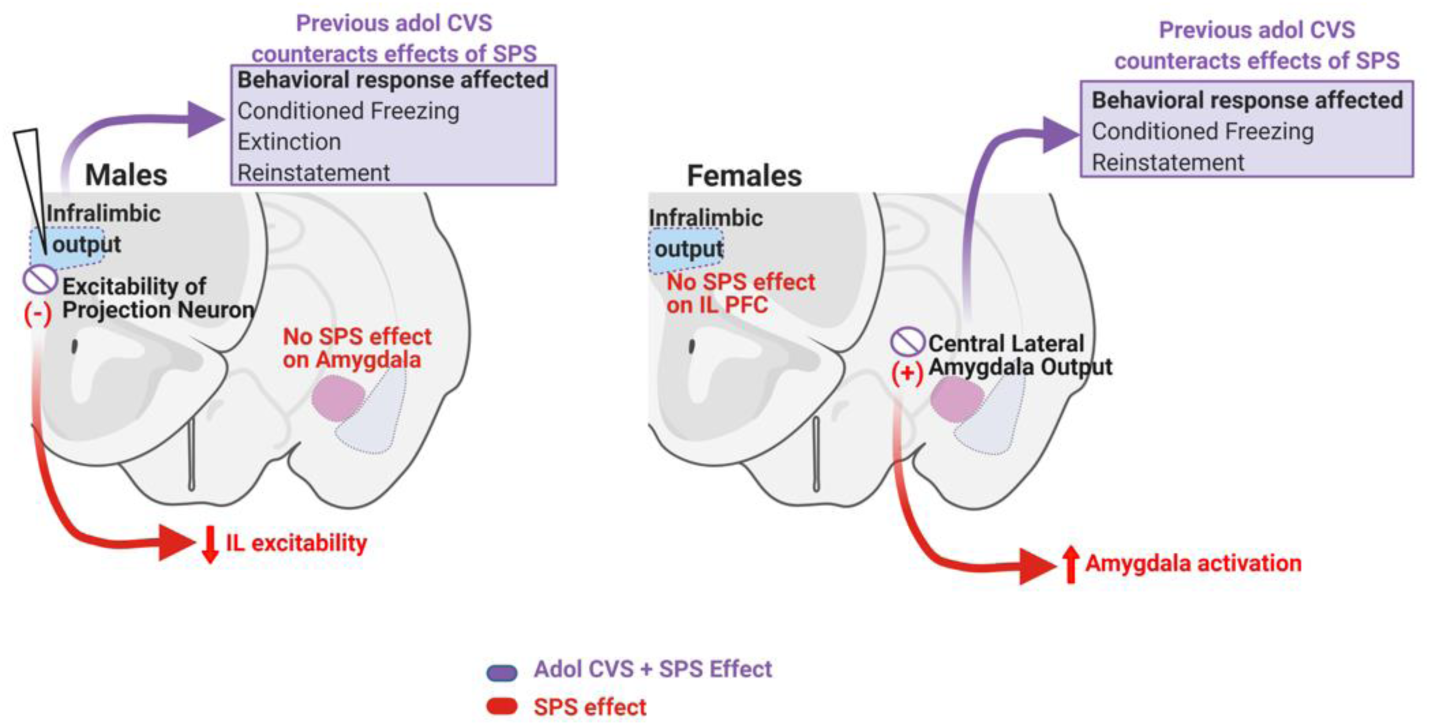
Summary of the effects observed. Prior exposure to adol CVS prevented the behavioral outcomes evoked by SPS in male and female rats. In the case of males, it also counteracted the effects of SPS on infralimbic excitability, indirectly inferred by Fos staining after reinstatement and confirmed by electrophysiology on pyramidal neurons from layer V. These central effects of adol CVS on SPS outcomes highlight the relevance of the infralimbic cortex as a hub for neurodevelopmental plasticity that could lead to resilience to stressful events in adulthood. In female rats, the absence of SPS effects in neuronal recruitment in the infralimbic cortex, accompanied by an increase of it in the central lateral division of the amygdala suggests a marked difference in the circuitry affected by SPS in both sexes. This points out to a possible mechanism involving excitability in that area that has yet to be confirmed.

Stress during development is generally thought to evoke negative behavioral effects later in life (Begni et al., 2020; Bourke and Neigh, 2011; Cotella et al., 2020, 2019; Green et al., 2013; Negrón-Oyarzo et al., 2014; Wilkin et al., 2012; Wulsin et al., 2016). However, prior studies also support the ability of adolescent stress to confer stress resilience in adulthood, using a number of stress models, e.g., intermittent predator stress (Kendig et al., 2011) and predictable chronic mild stress (PCMS) (Suo et al., 2013). Adolescent PCMS enhances extinction and prevents reinstatement and spontaneous recovery in a fear conditioning model evaluated immediately and one week following PCMS (Deng et al., 2017). Consistent with our results, these suggest that adolescent stress enhancement of resilience endures well beyond the time of exposure. The impact of adolescent stress differs thatt of stress imposition earlier in life, where the data generally report detrimental effects of stress (Johnson and Casey, 2014; Lukkes et al., 2009; McEwen, 2007; Vyas et al., 2002; Yee et al., 2012).

Although some authors proposed that the resilient phenotype is promoted by the predictability of the stressors (Deng et al., 2017), the general unpredictable nature of CVS suggests that the resilience mechanism is independent of response habituation In our study, the adol CVS paradigm employs exposure to swim and restraint, albeit in an isolated and time-attenuated fashion relative to SPS. Nonetheless, the length and consecutive application of the stressors during SPS represents a distinct and intense unpredictable experience. This contention is supported by a recent report demonstrating behavioral resilience to SPS using exposure to completely different stressors during adolescence (Chaby et al., 2020).

Timing combined with stressor modality seem to be an important factor as well. In this sense, prior work indicates adult resilience even after a single intense stressor protocol at PND37 (Moore et al., 2014) or following 3 days of predator related stressors at PND33-35 (Chaby et al., 2020). In contrast, a 3-day pre-pubertal exposure to variate stressors failed to attenuate exaggeration of fear responses in adulthood (Tsoory et al., 2010; Yee et al., 2012), indicating that developmental timing is critical for establishment of resilience.

Hypoactivity of the medial PFC is observed in several mental health disorders, including PTSD (Hains and Arnsten, 2008). Results from our group and others indicate that stress during adolescence reduces neuronal recruitment (Fos expression) to adult stressors in the mPFC (Cotella et al., 2020, 2019; Ishikawa et al., 2015). In humans, PTSD has been associated with a reduction in prefrontal drive, leading to abnormal extinction of conditioned fear (Milad et al., 2009, 2006, 2005; Rauch et al., 2006). Similarly, reduced IL mPFC activity following SPS in male rats may underlie abnormal extinction of fear responses (Piggott et al., 2019). Consistent with these data, our results indicate reduced IL recruitment following SPS in male rats accompanied by higher freezing during extinction and reinstatement. The reduced engagement of the IL in response to conditioned cues during the reinstatement procedure in the SPS male group also suggests a possible reduction of IL activity occurring during the prior extinction procedure, which would explain the impairment of extinction learning previously observed only in males rats.

Neurons in the mPFC are specifically activated during stressful situations and modulate their responses to subsequent exposure to the same stressor experience (Jackson and Moghaddam, 2006), thus playing a critical role in eliciting adaptive responses to aversive stimuli (Milad and Quirk, 2002). Modification of PFC responses to the same stimulus can be mediated through altered glutamatergic or dopaminergic drive onto the mPFC projection neurons (Bagley and Moghaddam, 1997; Jackson and Moghaddam, 2004). Adolescent social defeat decreases adult NMDA receptor expression in the IL PFC, and also reduces freezing to fear conditioning (Novick et al., 2016). Thus, the enhanced excitability we observed in SPS rats with prior history of adol CVS might be a long-term adaptation to the reduced excitatory drive that may occur following adol CVS.

Intrinsic membrane properties play an important role indetermining the prefrontal excitatory/inhibitory balance, as they directly shape neuronal output by influencing the probability of a neuron firing an action potential in response to synaptic inputs (Anderson et al., 2019). Our data indicate that the IL intrinsic excitability changes do not manifest at baseline conditions under adol CVS alone, consistent with prior work in mice resilient to social defeat (Friedman et al., 2014; Han and Nestler, 2017). Thus it is possible that prior adol CVS may serve to prime the pyramidal cells to react appropriately when faced with the second hit of SPS, compensating for reduced excitability associated with SPS. The exact mechanism underlying the altered excitability of IL pyramidal neurons observed in our study is yet to be determined. Possibilities include lasting alteration in ion channel function (e.g., G protein-gated inwardly rectifying K+ channels) (Anderson et al., 2019; Hearing et al., 2013) or modulation on excitability by hyperpolarization-activated cyclic nucleotide–gated channels (Shah, 2014). Further work is needed to identify the specific ionic mechanisms by which adolescent stress can protect against future stressors during adulthood. (Matovic et al., 2020).

## Conclusion

Our results support the idea that certain combinations of stressful situations during adolescence can be beneficial, evoking resilience to stress in adult life. We propose that, in rats, chronic variable stress during late adolescence determines differential activation or recruitment of the IL in response to intense stress in adulthood. This rearrangement of prefrontal activity results in a phenotype that is resilient to stress-enhanced fear learning, reducing contextual response, facilitating extinction and preventing reinstatement of the fear conditioned response following trauma, findings that may lend insight into under standing susceptibility of resilience toPTSD. Furthermore, our data guide our next steps to understand the sex specific effects in behavioral resilience following adolescent stress that pointed to fundamental sex differences in stress reactive brain regions and their involvement in resilienceIt would be important as well to determine which stressor type might result in a positive emotional valence and whether that ultimately evokes a resilience response. For example, the exercise (swim) or social component (crowding) of the adolescent CVS regimen might help individuals cope with subsequent stress in adulthood (Herring et al., 2010; Ozbay et al., 2007). The next challenge is to find the most efficient developmental triggers for the generation of resilience to the effects of adult stress, possibly including positive developmental interventions, with the goal of reducing the incidence of stress-related affect conditions, including PTSD.

## Materials and Methods

### Animals

Male and female Sprague Dawley rats were bred in-house, weaned at postnatal day 21 (PND21) and pair-housed in standard clear cages (20 cm height x22 cm width x43 cm length) under a 12 h light/ 12 h dark cycle (lights on at 7:00 am), at constant room temperature (23+2°C), with *ad libitum* access to food and water. All tests were performed during the light cycle, between 09:00 AM and 2:00 PM. All procedures and care performed in the animals were approved by the University of Cincinnati Institutional Animal Care and Use Committee.

### Adolescent chronic variable stress (adol CVS)

After weaning the animals, a cohort was randomly assigned to 4 experimental groups Control (No CVS – No SPS), Adol CVS (only chronic stress in adolescence), SPS (only single prolonged stress in adulthood) and CVS-SPS or double-hit group (adol CVS + SPS in adulthood). The remaining animals from the litter were assigned to control and adol CVS groups for the evaluation in the novel object test and Morris water maze to assess general cognitive function. No more than 2 littermates were included in each experimental group. Rats were subjected to our standard 14 days CVS protocol during late adolescence (PND 45 ± 2) following prior work from our group (Jankord et al., 2011). The **<u>CVS paradigm</u>** included a set of unpredictable variable stressors applied twice daily (AM and PM, see table). In addition, animals were exposed to overnight stressors every two days: 1) individual housing, 2) social crowding (six rats per cage). Control animals were maintained in the same room and only handled for normal husbandry. Except for overnight stressors, all the procedures related to CVS were performed in a different room. Following CVS, animals were allowed to recover for 4 weeks to be then subjected to the single prolonged stress (SPS) protocol evaluated during adulthood. The timeline of the experiment is shown in Fig. 1.

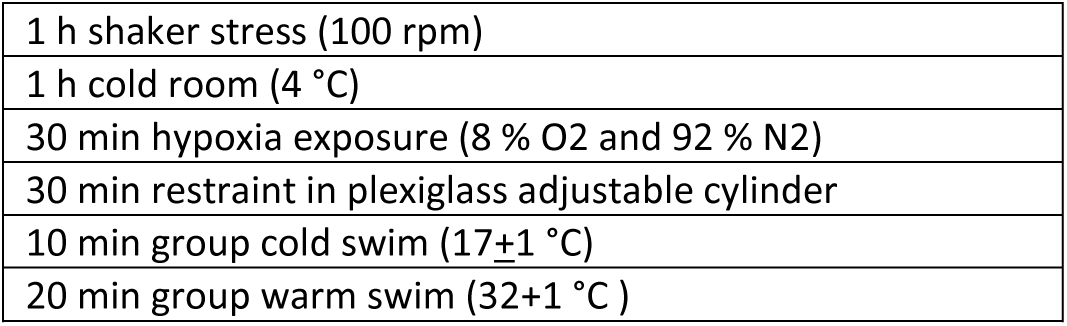

### Single prolonged stress (SPS)

A groups of rats subjected to adolescent CVS (and their respective controls) were subjected to SPS 4 weeks after the end of CVS. For this, starting at 9:00 AM, animals were restrained for 2 hours in plexiglass adjustable cylinders dimensions 20cm X 7 cm. Immediately after 2h elapsed, they were subjected to 20 minutes of group swim (25 + 2°C) in a bucket with dimensions 50cm X 33cm. Immediately after they were retrieved from the water and the excess of water was eliminated from their coat, they were allowed to recover for 15 minutes in their home cage with their cage mate. Next, rats were placed into a glass chamber where they were exposed to ether vapor until loss of consciousness (loss of righting position and palpebral reflex). Immediately after unconsciousness was confirmed they were placed in a cage with clean bedding material and returned to their housing room. SPS was administered in a novel experimental room. After SPS, animals remained undisturbed for a week (the usual time required to observe SPS effects (Knox et al., 2012b; Kohda et al., 2007; Wen et al., 2015)). Control rats remained in the housing room during the application of SPS and were only subjected to cage change during that time.

### Cued Fear Conditioning Paradigm

A week after SPS all groups were subjected to an auditory tone cued fear conditioning protocol to evaluate the performance of the animals during the conditioning, extinction, and reinstatement sessions. Behavioral evaluation occurred between 8:30 AM – 2:00 PM. **<u>Conditioning:</u>** animals were allowed to explore the conditioning chamber for 3 min, after which they were exposed to a 20s auditory tone (conditioned stimulus, CS), co-terminating with a 0.5s, 0.45mA shock (unconditioned stimulus, US), with an inter-trial interval (ITI) of 120s. The tone-shock pairings were repeated six times. Data were presented as % Freezing over CS time (20s). **<u>Extinction:</u>** 24 h after conditioning animals were subjected to 3 consecutive days of extinction in which they were exposed to 7 repetitions of the CS with 120s ITI in the same conditioning chambers. Data were presented as Total % Freezing over the 3 days of the extinction procedure, corresponding to the cumulative % of freezing expressed over the cumulative CS time during the whole extinction session (7×20s: 140s). **<u>Recall test:</u>** 5 days after the last extinction session animals were placed in the conditioning chambers and exposed to 3 repetitions of the CS, with 120 s ITI with the purpose of evaluating spontaneous recovery of the conditioned response. Data were presented as Total % Freezing calculated as cumulative % of freezing expressed over the cumulative CS time during the session (3×20s: 60s).

### Reinstatement

48h after the recall session animals were exposed to a reinstatement session in the same conditioning chambers, consisting of 3 min of chamber exploration followed by 1 unpaired shock (US) (0.45 mA, 0.5s). After a delay of 120s animals were exposed to 5 repetitions of the CS (120s ITI). Data were expressed as total % Freezing during tones after unpaired shock, Total % Freezing over the cumulative duration of CS time (5×20s= 100s). Rats were euthanized 90 min after the onset of the session to obtain brains for immunoshitochemistry.

### Context fear conditioning test

As a way of quantifying the conditioned response to the context, we evaluated the initial freezing response exerted every day of the extinction procedure before the first tone was presented and analyzed the progression of this response over the 3 days. Data were expressed as % Freezing over the initial pre-tone exploration time (120s).

The conditioned response evaluated was freezing behavior, considered as general absence of movement, which was scored using a video tracking system (EthovisionXT-Noldus). We did not consider other behaviors as, in our set up, animals do not show conditioned darting or any other escape related behaviors, with both sexes consistently exhibiting freezing in response to the tone. 90 minutes after reinstatement rats were euthanized to obtain their brains for immunodetection of Fos, a marker of neuronal activation. Results were analyzed considering sex as a variable.

Group composition fear conditioning experiment: The experiment was originally designed with 10 male rats per group and 12 female rats per group. A male rat assigned to CVS died of unknown reasons during the first week of CVS resulting in a n of 9 for that group. A spare rat was added to that home cage to avoid having to exclude the house mate of the lost individual due to isolation. Later, on extinction day 2, there was a malfunction of the conditioning chambers which resulted in shocking the animals as soon as the session started. This resulted in the exclusion of the 3 animals run during that session (2 controls, 1 CVS). Therefore beginning day 2 of the paradigm, the n for control and CVS males was 8.

In the case of females, we originally planned to include 12 females per group, nevertheless, due to a mistake during the day of SPS, 1 cage of CVS animals was wrongfully submitted to SPS in lieu of a cage without CVS planned to get SPS. That resulted in an imbalance in the final number of animals per group: Control and CVS-SPS groups had n=14 and CVS and SPS groups had n=10. The experimental timeline is shown in Figure 1.

### Immunohistochemistry

Rats were euthanized with an overdose of sodium pentobarbital and immediately transcardially perfused with 0.9 % saline followed by 4 % paraformaldehyde in 0.1M phosphate buffer (PBS), pH 7.4. Brains were post-fixed in 4 % paraformaldehyde at 4 °C for 24 h, then transferred to 30 % sucrose in 0.1 M PBS at 4 °C where they were kept until tissue processing. Brains were sliced into serial 35 μm coronal sections using a freezing microtome (−20 °C). Sections were collected into multi-well plates containing cryoprotectant solution (30 % Sucrose, 1 % polyvinyl-pyrolidone (PVP-40), and 30 % ethylene glycol in 0.1M PBS). For immunolabeling, sections were washed 6×5 min in 0.01M PBS at room temperature (RT). After being rinsed, sections were incubated with 1 % sodium borohydride in 0.1 M PBS for 30 min at RT. After rinsing 6×5 min 0.1 M PBS, they were incubated in 3 % hydrogen peroxide diluted in 0.1M PBS for 20 min. Subsequently, brain slices were rinsed 6×5 min and 4×15 min in 0.1M PBS and then incubated in blocking solution (4 % normal goat serum (NGS), 0.4 % TritonX-100, 0.2 % bovine serum albumin (BSA) in 0.1M PBS, 2 h at RT. Sections were then incubated with c-Fos rabbit polyclonal antibody (1:1000, Santa Cruz, sc-52) in blocking solution, overnight at RT. The next day, sections were rinsed (3×5 min) in 0.1 M PBS at RT, followed by incubation with secondary antibody (biotinylated goat anti-rabbit, 1:400; Vector Laboratories, BA1000) in blocking solution at RT for 1 h. Sections were again rinsed (3×5 min) in 0.1 M PBS and then reacted with avidin-biotin horseradish peroxidase complex (1:800 in 0.1 M PBS; Vector Laboratories) for 1 h at RT. Sections were then rinsed (3×5 min) in 0.1 M PBS and then developed with a 8 min incubation in DAB-Nickel solution: 10 mg 3,3′-diaminobenzidine (DAB) tablet (Sigma, DF905), 0.5 ml of a 2 % aqueous nickel sulfate solution, 20ul of 30 % hydrogen peroxide in 50 ml of 0.1 M PBS. Sections were finally washed in PBS, mounted on superfrost slides (Fisherbrand, Fisher), allowed to dry, dehydrated in xylene, and then coverslipped in DPX mounting medium (Sigma). Sections from 5 to 7 brains per experimental group were processed. For analysis, we counted 3 bilateral sections from equivalent coordinates covering the anterior, medial and posterior portions of the prefrontal cortex (PFC), nuclei in the amygdala: central amygdaloid nucleus (CeAm), medial amygdaloid nucleus (MeAm), lateral amygdaloid nucleus (LAm) and basolateral amygdala (BLA). Each brain region limit and coordinates were defined following a brain atlas (Paxinos and Watson, 2007). The number of Fos positive nuclei was counted with a semiautomatized method using ImageJ software (National Institutes of Health, Bethesda, MD). Counts of Fos immunoreactive cells were obtained from each area of interest using the Analyze Particle tool, using a defined common level of background intensity, nuclei circularity and size (previously validated manually). Once the number of Fos positive nuclei was determined in each section, the relative density of the population of immunopositive cells was calculated by dividing this number by the area measured in each case. Considering that the number of animals used simultaneously in the study makes it logistically complicated to process all the tissue at the same time or separating in batches and obtain homogenous immune staining, we decided to prioritize the within sex results for the Fos quantification and tissue from male and female rats was processed and analyzed independently.

### Electrophysiology

Following the same timeline as the behavioral studies, whole-cell patch clamp recordings were obtained from layer V pyramidal neurons in the IL PFC. Details of slice preparation and electrophysiology recordings from adult PFC are given below.

### Slice Preparation

Rats were sacrificed 7 days post SPS. Animals were deeply anesthetized with sodium pentobarbital (390 mg/kg, Fatal-Plus) and decapitated. A warm slicing protocol was used to prepare healthy adult rat brain slices as previously described (Ting et al., 2014). Brains were quickly isolated and dura matter carefully removed before removing the cerebellum. The brain was then immediately glued to a cutting stage immersed in NMDG solution (92 mM NMDG, 2.5 mM KCl, 1.2 mM NaH_2_PO_4_, 30 mM NaHCO_3_, 20 mM HEPES, 25 mM glucose, 5 mM sodium ascorbate, 2 mM thiourea, 3 mM sodium pyruvate, 10 mM MgSO_4_, and 0.5 mM CaCl_2_) at a temperature of 34-36°C and continuously bubbled with 95% oxygen and 5% carbon-dioxide. Coronal slices containing the mPFC were sectioned at 300 µm thickness using a vibrating microtome (7000smz-2; Campden Instruments, Lafayette, IN) with ceramic blades (Campden Instruments) at an advance speed of 0.03 mm/s. Vertical vibration of the blade was manually tuned in accordance with the user manual, and was set to 0.1 – 0.3 μm. Bath temperature was kept within the desired range of 34-36°C, by adding warm or cold water into the external chamber of the slicer, and was monitored throughout the cutting procedure with a conventional mercury/glass thermometer. The slices were allowed to recover for 1 hour in oxygenated NMDG solution at 34-36°C. At the end of recovery, slices were transferred to a chamber containing oxygenated artificial CSF solution (125 mM NaCl, 2.5 mM KCl, 25 mM NaHCO_3_, 1 mM NaH_2_PO_4_, 25 mM glucose, 1 mM MgCl_2_, 2 mM CaCl_2_) for at least 30 minutes at room temperature after which the slices were ready for in vitro patch clamp recordings for the next 1-6 hours.

### Electrophysiological recording

Brain slices were transferred to a submersion-type recording chamber (RC-22; Warner Instruments, Hamden, CT) and mounted onto the stage of an upright microscope (BX51WI, Olympus, Center Valley, PA). Slices were then perfused at a flow rate of 2–4 ml/min with oxygenated aCSF at 34-36°C. Patch electrodes were constructed from thin-walled single-filamented borosilicate glass (1.5 mm outer diameter; World Precision Instruments) using a microelectrode puller (P-97; Sutter Instruments, Novato, CA) and filled with an intracellular solution (130 mM K-gluconate, 10 mM KCl, 10 mM HEPES, 10 mM sodium phosphocreatine, 4 mM MgATP, and 0.3 mM Na_2_-GTP, pH 7.2, 295-300 mOsm). Pipette resistances ranged from 4 to 6 MΩ, and seal resistances were > 1 GΩ.

Whole-cell patch clamp recordings were obtained from layer V pyramidal in the mPFC using a MultiClamp 700B Amplifier (Molecular Devices, Sunnyvale, CA). Pyramidal neurons were easily identifiable in the slice based on soma morphology and the presence of a prominent apical dendrite. In the current clamp mode, once a stable membrane potential was observed, intrinsic excitability measurements were performed at the resting membrane potential (RMP). Cell capacitance was measured using the membrane test function in pClamp 10.4 (Molecular Devices, Sunnyvale, CA, USA). All measurements of intrinsic membrane excitability were taken from RMP. Rheobase was measured by applying depolarizing current steps (10 pA steps, 100 msec duration) until the generation of a single action potential (AP). Input resistance was measured by applying a hyperpolarizing current step (−20 pA) via the patch electrode. AP threshold was defined as the *V*m measured 0.5ms before the peak in the second derivative of the waveform. The action potential threshold and amplitude were analyzed for the first spike at the rheobase current injection. Duration of APs (AP_50_) was determined by measuring the elapsed time from the peak of the AP to 50% maximum amplitude during the repolarization phase. Cells with RMP lower than -55mV were included for the final analysis. Outliers were detected using the Grubbs’ test (GraphPad Software) and removed from analysis. For AP_50_: 1 neuron from control and adol CVS+ SPS group; RMP: 1 neuron from control and adol CVS; AP amplitude: 1 neuron from adol CVS group; Membrane resistance: 1 neuron from adol CVS + SPS group; Capacitance: 1 neuron from control and 3 neurons from adol CVS group and for AP threshold 3 neurons from adol CVS and adol CVS+SPS were removed as outliers. Firing rate was measured in response to 20 pA and 1 sec duration depolarizing current steps in the current clamp configuration. Only cells that produced greater or equal to 5 action potentials for up to 240 pA current injection were included in the final analysis. Peak firing frequency was reported as the maximum number of action potentials generated in a given neuron following a current injection step. Number of cells used for electrophysiology are outlined in table 1. Membrane voltages were adjusted for liquid junction potentials (approximately –14 mV) calculated using JPCalc software (P. Barry, University of New South Wales, Sydney, Australia; modified for Molecular Devices). Signals were filtered at 4–6 kHz through a –3 dB, four-pole low-pass Bessel filter and digitally sampled at 20 kHz using a commercially available data acquisition system (Digidata 1550A with pClamp 10.4 software). Data were recorded using pClamp, version 10.4 (Molecular Devices) and stored on a computer for offline analysis. Recordings were detected and analysed using Clampfit (Molecular Devices).

**Table 1.** Table depicts number of cells used for the electrophysiology data analysis

|  | Number of cells |
| --- | --- |
| a | Rheobase was measured in 19 neurons in the control group, 17 neurons in SPS group, 35 neurons in adolescent stress group, 31 neurons in adolescent stress and SPS group n=2-3 animals/group |
| b | Membrane resistance was measured in 19 neurons in control group, 17 neurons in SPS group, 35 neurons in adolescent stress group and 30 neurons in the adolescent stress and SPS group n=2-3 animals/group |
| c | AP threshold was measured in 19 neurons in control group, 17 neurons in SPS group, 32 neurons in adolescent stress group and 28 neurons in the adolescent stress and SPS group n=2-3 animals/group |
| d | AP amplitude was measured in 19 neurons in control group, 17 neurons in SPS group, 34 neurons in adolescent stress group and 31 neurons adolescent stress and SPS group n=2-3 animals/group |
| e | AP<sub>50</sub> was measured in 18 neurons in control group, 17 neurons in SPS group, 35 neurons in adolescent stress group and 30 neurons in the adolescent stress and SPS group n=2-3 animals/group |
| f | Resting membrane potential was measured in 18 neurons in control group, 17 neurons in SPS group, 34 neurons in adolescent stress group and 31 neurons in the adolescent stress and SPS group n=2-3 animals/group |
| g | Capacitance was measured in 19 neurons in control group, 17 neurons in SPS group, 32 neurons in adolescent stress group and 31 neurons in the adolescent stress and SPS group n=2-3 animals/group |
| h | Peak firing frequency was measured in 11 neurons in control group, 8 neurons in SPS group, 5 neurons in adolescent stress group and 16 neurons in the adolescent stress and SPS group n=2-3 animals/group |

## Statistical analysis

Fear conditioning data were analyzed by repeated measurements ANOVA (adol CVS x SPS x Sex x time), with a level of significance of p < 0.05. Novel object recognition and Morris water maze data were analyzed by 2×2 ANOVA (Adol CVS x Sex). Fos data were analyzed by a 2-way ANOVA (2×2 design: adol CVS x SPS) within each sex with a level of significance p < 0.05. Electrophysiology data were analyzed by 2×2 ANOVA (adol CVS x SPS). Details of number of cells used for electrophysiology data analysis are outlined in table 1. In the cases where significant differences and interactions were found, the Bonferroni test was used for post hoc analysis. In the case there were only main effects of the factors but no significant interaction between them, we performed planned comparisons to evaluate individual differences. Data were analyzed using STATISTICA 7.0 (Statsoft, Inc.,Tulsa, USA) and Prism 8 (GraphPad Software, La Jolla California USA). Data not following a normal distribution were log transformed for statistical analysis.

## Acknowledgements

The authors would like to thank other members of Dr Herman’s laboratory for their assistance in data collection and general discussion of the results. Images were created with BioRender.com.

